# Learning Shapes the Energy Cost of Neural Tasks

**DOI:** 10.64898/2026.07.01.735889

**Authors:** Kaili Xue, Farid Rezayat, Tianbo Qi, Leyao Shen, Bohan Zhao, Xun Huang, Jonathan S. Marvin, Li Ye

## Abstract

The stark difference in energy use between AI systems and biological brains highlights both the remarkable efficiency of the brain and our limited understanding of the energetics underlying neural computation. Although efficiency is widely cited as a principle of neural design, direct measurement of the energy cost of specific neural tasks within their corresponding circuits has remained limited. Here, by simultaneously measuring intracellular glucose and calcium dynamics in the same behaving mice *in vivo*, we use neuronal glucose consumption to define circuit-level energy costs associated with learning-based behavioral tasks. We found that the post-learning fuel cost per task was significantly lower than pre-learning levels across multiple hippocampus- and cortex-dependent learning models, with or without changes in bulk calcium dynamics. This change in fuel cost was not mediated by extracellular glucose transport but instead reflected a reduction in intracellular glucose consumption and depended on canonical plasticity mechanisms, including NMDAR signaling and protein synthesis. Together, these findings suggest that attenuation of task-specific energy cost may represent a general bioenergetic trajectory of learning and plasticity. This “energy minimization” hypothesis provides insight into how the biological brain achieves efficiency and offers an orthogonal, complementary perspective to neural activity-centered frameworks.

## Main Text

In contrast to the gigawatt scale energy consumption of AI, the biological brain operates at incredibly low power. For example, it is estimated that the human brain operates on a modest, nearly constant 20 W of power, even when performing complex tasks or learning. The striking energetic difference between these two neural networks highlights a fundamental gap in our understanding of the biological brain. Moreover, since the brain secures and regulates its own power supply by controlling organismal physiology and behavior, it must use efficient strategies to obtain, store, and, most importantly, spend this precious energy.

Energy efficiency has been recognized as a general principle of neural design (*1–4*) and a driving selection force of brain evolution (*5*); for example, many theoretical models suggest metabolically efficient wiring patterns (*6, 7*) and sparse coding (*8–10*) lower the energy cost of neural computation. However, their integration into modern neuroscience frameworks remains limited. Most brain models treat metabolic cost as a fixed background assumption rather than a dynamic variable, leaving the dynamic relationship between energy expenditure and learning largely underspecified. Although “efficiency” is widely cited as a driver of neural principles in ethology, this has been difficult to experimentally quantify in a functioning neural circuit. Neuroimaging methods like BOLD/fMRI and FDG-PET are largely based on local brain energy metabolism, but they are often used as an approximation of neural activity, rather than to be used as a measure of the energy cost of neural computation (*11–15*). Thus, to date, most estimates of the energy cost of information processing are based on theoretical models rather than direct experimental measurement, precluding a complete understanding of brain energetics.

Glucose is the primary energy source of the brain under normal physiological conditions, and activity-dependent changes in neuronal glucose provide a physiologically grounded readout of local energetic demand (*16–19*). Conventional glucose measurements based on enzymatic assays or radioactive imaging, however, lack either the speed to match the sub-second temporal resolution of neural dynamics *in vivo* or the spatial and cellular specificity to resolve circuit-level glucose changes. The recent development of genetically encoded fluorescent glucose sensors, iGlucoSnFR (*20*) and its improved version iGlucoSnFR2 (*21*), enables sub-second temporal resolution to track glucose changes in extracellular and intracellular compartments *in vivo* and in a cell type specific manner (*21*).

By combining iGlucoSnFR2 with in brain fiber photometry in parallel with calcium imaging, we monitored neural activity-linked intracellular glucose consumption in freely moving, behaving mice. The high spatial and temporal resolution of this approach also allowed us to use the rate of intracellular glucose consumption as a surrogate to estimate circuit-selective energy costs associated with behavior-defined neural tasks. We found that task-specific energy consumption rises sharply during the initial learning phase but falls as the task becomes routine. This post-learning reduction in energy consumption is consistently observed in multiple well-established learning-based behavioral paradigms in the hippocampus and cortex and across various time scales. Based on these findings, we propose an “energy minimization hypothesis” that holds that reducing the energy cost of a neural task is a general bioenergetic outcome of learning and can be measured orthogonally to neural activity itself.

### Visual activity transiently lowers intracellular glucose in primary visual cortex neurons

We first used well-established visual stimulation-induced neural activity in primary visual cortex (V1) neurons as a model to test how sensory-induced activity affects neuronal glucose dynamics. To monitor glucose in distinct cellular compartments, we expressed cytosolic and extracellular membrane-targeted configurations of iGlucoSnFR2 under the hSyn promoter in neurons on the left and right sides of V1, respectively, as previously reported (*21*) (Fig. 1, A and B). iGlucoSnFR2 provides improved sensitivity and reliability compared with iGlucoSnFR1, enabling *in vivo* monitoring of cytosolic glucose dynamics in the mammalian brain (*21*). We confirmed that intraperitoneal glucose injection leads to an increase in both intracellular and extracellular glucose in V1 neurons (Fig. 1, C and D), indicating both configurations of the iGlucoSnFR2 sensor were functional.

**Fig. 1.**
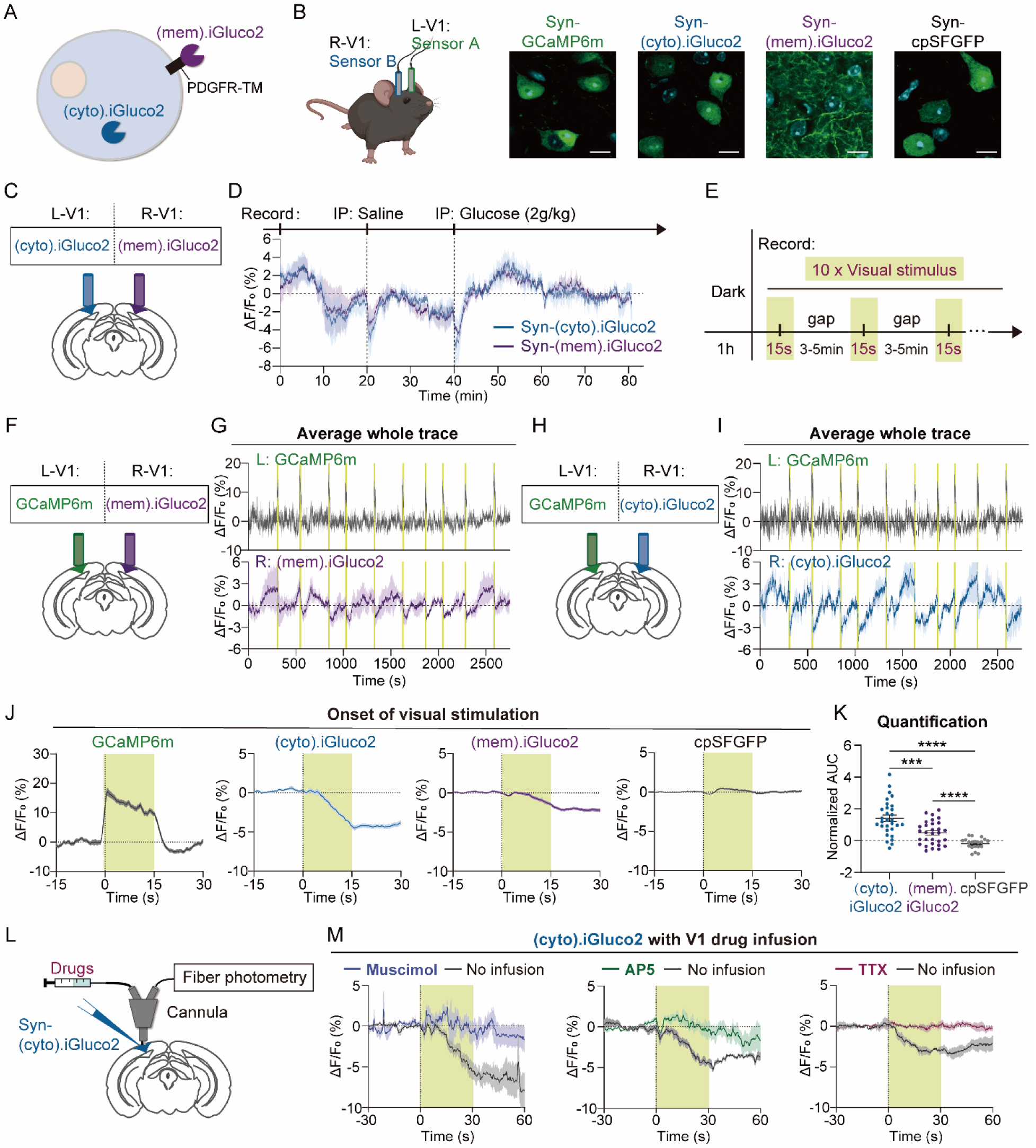
Visual activity transiently lowers intracellular glucose in primary visual cortex neurons. (A) Schematic of the two iGlucoSnFR2 sensor configurations: subcellular targeting to the cytosol (blue, (cyto).iGluco2) and to the plasma membrane (purple, (mem).iGluco2). PDGFR-TM, Platelet-derived growth factor receptor transmembrane domain. (B) Schematic of AAV injection and fiber placement (left). AAVs encoding sensors are injected into the left and right V1 in the same animal (as illustrated in C, F, and H). Histological verification is shown in the right panel (scale bar, 20 μm). (C) Schematic showing sensor injection and fiber placement for fiber photometry recording from V1 neurons with intraperitoneal glucose injection for functional validation of the (cyto).iGluco2 and (mem).iGluco2 sensors. (D) Average fiber photometry traces showing (cyto).iGluco2 (blue) and (mem).iGluco2 (purple) signal dynamics in response to systemic saline and glucose injection (n = 3 mice). (E) Visual stimulation paradigm: mice were first habituated in the dark for 1 hour, followed by ten 15-second visual stimulation trials during fiber photometry recording. (F) Schematics showing sensor injection and fiber placement for visual stimulation experiments in G. (G) Average fiber photometry traces showing GCaMP6m signals in the left V1 (upper) and (mem).iGluco2 signals in the right V1 (lower) of the same animal during visual stimulation (n = 3 mice). (H) Schematics showing sensor injection and fiber placement for visual stimulation experiments in I. (I) Average fiber photometry traces showing GCaMP6m signals in the left V1 (upper) and (cyto).iGluco2 signals in the right V1 (lower) of the same animal during visual stimulation (n = 3 mice). (J) Signal changes aligned to the onset of visual stimulation (n = 3 mice). (K) Quantification of (cyto).iGluco2, (mem).iGluco2, and cpSFGFP signals shown in J. Normalized area under the curve (AUC) was calculated over the 15-s visual stimulation period and compared across groups. (L) Schematic of sensor injection and cannula placement for local drug infusion in V1 during fiber photometry recording. (M) (cyto).iGluco2 signal changes aligned to visual stimulation onset with and without local drug infusion. A GABA_A_ receptor agonist (muscimol, 0.125 μg/site), an NMDAR antagonist (AP5, 0.1 μg/site), or a sodium channel blocker (TTX, 2 ng/site) was infused into V1 through the cannula. Statistics were determined using an unpaired t test in (K). *P < 0.05; **P *<* 0.01; ***P< 0.001; ****P<0.0001.

For sensory stimulation, mice were first habituated in the dark for 1 hour, then exposed to a set of visual stimulus (Fig. 1E). GCaMP6m was expressed in neurons on the left side of the V1 while intra- or extracellular iGlucoSnFR2 was expressed on the right side of the V1 in the same animals (Fig. 1, F and H). Visual stimulation led to time-locked neural activity, as indicated by GCaMP6m peaks (Fig. 1G and fig. S1A). As expected, given the coupling between neurons and the vasculature, this transient activity caused a rapid drop of extracellular glucose which quickly recovered after visual stimulation (Fig. 1G and fig. S1A). We reasoned that the extracellular glucose dynamics *in vivo* (e.g., neuron-vasculature coupling) would be able to maintain intracellular glucose, at least at the soma, at a stable level (*22–24*). However, unexpectedly we observed an even steeper drop in intracellular glucose levels following visual stimulation (Fig. 1, I to K, and fig. S1B). This difference between extra- and intra-cellular glucose was more apparent when the visual stimulus was extended to 30 minutes (fig. S1, D to F), where extracellular glucose recovered much faster than the intracellular level. To rule out the possibility that this drop was due to changes in intracellular pH or other non-glucose induced iGluco signals, we used a sensor-dead cpSFGFP, commonly used to control fluorescence sensors (*25*), under the same behavioral paradigm, but cpSFGFP did not display similar visual stimulation-coupled signal decreases (Fig. 1, J and K, and fig. S1C). Moreover, the iGlucoSnFR2 readout of intracellular glucose could be lowered by local infusion of a pan-class I GLUT (GLUT 1-4) transporter inhibitor, DRB18 (*26*) (fig. S2A), suggesting iGlucoSnFR2 is reporting a bona fide glucose drop upon visual stimulation. We also observed a similar, but less steep drop of intracellular glucose in astrocytes (fig. S2, C to E) under the same experimental conditions.

Neural activity is known to increase intracellular glucose metabolism (*27, 28*). Thus, this apparent drop in intracellular glucose level likely reflects a transient imbalance in which neural activity-induced glucose consumption exceeds glucose influx. This is supported by the further amplification of the visual stimulation-induced decrease in glucose following local infusion of the GLUT inhibitor DRB18 (fig. S2B). To test if neural activity is driving this drop, we repeated the visual stimulation paradigm in the presence of different classes of pharmacological blockers of neuronal activity, including a GABAR agonist (muscimol), an NMDAR antagonist (AP5), or a Na channel block (TTX) (fig. S2F). We found that these drugs effectively abolished the VS-induced intracellular glucose drop (Fig. 1, L and M), suggesting that neuronal activity drives glucose consumption inside the neurons.

### Learning shapes glucose consumption of CA1 neurons during spatial navigation

In our routine experiments, we noticed that repeated visual stimulation on the same individual resulted in diminished intracellular glucose drops in V1 neurons, suggesting that learning and anticipation may influence neuronal glucose consumption. To formally test how learning may affect neural glucose consumption, we measured glucose levels while mice performed a well-established maze navigation task in which spatial learning can be readily tracked by behavioral quantification.

In this paradigm, we expressed GCaMP6m in CA1 neurons on one side of the hippocampus and iGlucoSnFR2 on the contralateral CA1 in the same mouse (Fig. 2, A and B). Mice were trained to navigate a modified Hebb–Williams-type maze (*29*) for a water reward (Fig. 2C), and they quickly learned this task in 1-3 trials, as reflected by the significant reduction in time to reward from > 1 minute to < 20 seconds (Fig. 2D) and the acquisition of the shortest paths between the entry and reward (Fig. 2E). The first trial, with the highest “effort” and GCaMP6m activity, is associated with the sharpest intraneuronal glucose drop (Fig. 2, F to H, and fig. S3A). With subsequent trials, the navigation task consumes much less intracellular glucose as the mice navigate the maze more quickly and with lower bulk neural activity (Fig. 2, G, H, and O). This decrease of spatial navigation related bulk neural activity is consistent with classic electrophysiology studies showing that in the CA1, the average firing rate (FR) drops between the first and subsequent trials in a new maze (*30*). Our experimental measurements directly support the notion that such sharper, high SNR coding after learning translates to reduced energy expenditure to complete the same navigation task (*8*).

**Fig. 2.**
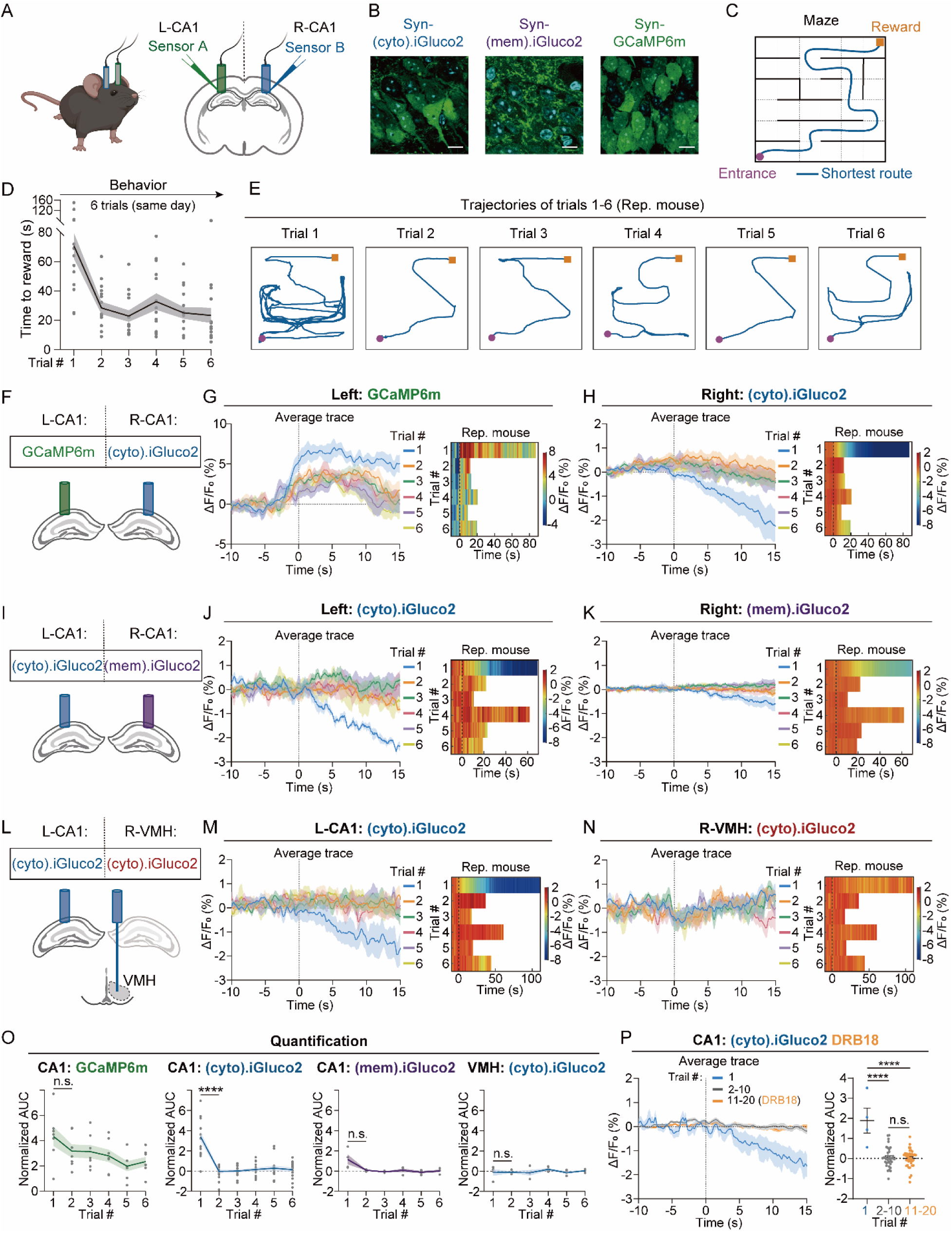
Learning shapes glucose consumption of CA1 neurons during spatial navigation. (A) Schematic showing sensor injection and fiber placement. (B) Histological verification of sensor expression in CA1 neurons (scale bar, 20 μm). (C) Schematic of the maze layout. Black lines indicate the maze walls, and the blue line marks the shortest path from the entrance to the reward. (D) Maze navigation learning curve showing time to reward across trials 1–6 (n = 15 mice). (E) Maze trajectories from one representative mouse across trials 1–6. (F) Schematic showing sensor injection and fiber placement for the experiments in G and H. (G, H) Trial onset-aligned average traces showing GCaMP6m (G) and (cyto).iGluco2 (H) signals in CA1 neurons across trials 1–6 from the same mice (left, n = 7 mice). Representative heatmaps from one mouse showing trial-by-trial GCaMP6m and (cyto).iGluco2 signals across trials 1–6 (right). (I) Schematic showing sensor injection and fiber placement for the experiments shown in J and K. (J, K) Trial onset-aligned average traces showing (cyto).iGluco2 (J) and (mem).iGluco2 (K) signals in CA1 neurons across trials 1–6 from the same mice (left, n = 4 mice). Representative heatmaps from one mouse showing trial-by-trial (cyto).iGluco2 and (mem).iGluco2 signals across trials 1–6 (right). (L) Schematic showing sensor injection and fiber placement for the experiments shown in M and N. (M, N) Trial onset-aligned average traces showing (cyto).iGluco2 signals in CA1 neurons (M) and VMH neurons (N) across trials 1–6 from the same mice (left, n = 4 mice). Representative heatmaps from one mouse showing trial-by-trial (cyto).iGluco2 signals in CA1 and VMH neurons across trials 1–6 (right). (O) Quantification of GCaMP6m, (cyto).iGluco2, and (mem).iGluco2 signals in CA1 neurons, as well as (cyto).iGluco2 signals in VMH neurons. The normalized AUC of individual trials 1–6 was quantified. (P) (cyto).iGluco2 signals in CA1 neurons with GLUT transporter inhibitor (DRB18, 10 ng/site) infusion during trials 11-20. The left panel shows the average signal traces for trial 1, trials 2–10, and trials 11–20 (DRB18) (n = 4 mice). The right panel shows the normalized AUC quantification for trial 1, trials 2–10, and trials 11–20 (DRB18). Statistics were determined using a paired t test in (O) and an unpaired t test in (P). *P < 0.05; **P *<* 0.01; ***P< 0.001; ****P<0.0001.

Following this post-learning reduction in intracellular glucose consumption, intracellular glucose levels stabilize and become insensitive to the inhibition of glucose import by DRB18 (Fig. 2P and fig. S4). This suggests that with tasks becoming routine, there is reduced neuronal glucose consumption rather than acute changes in fuel delivery or vascular transport. Indeed, in the same animals, extracellular glucose levels did not show post-learning reversals (Fig. 2, I to K and O, and fig. S3B). Moreover, intracellular glucose in the ventromedial hypothalamic nucleus (VMH), a center for regulating systemic glucose metabolism, did not exhibit post-learning changes in glucose levels during this spatial navigation task (Fig. 2, L to O, and fig. S3C). This indicates that the post-learning reduction is highly specific to the hippocampus, where spatial learning primarily happens, and not a reflection of systemic changes in glucose levels associated with locomotion or global metabolism.

### Glucose consumption rebounds during a renewed spatial navigation task

We next examined glucose consumption during error-driven spatial relearning by modifying a familiar maze, thereby requiring animals to remap the optimal route to the reward. Mice were familiarized with the initial maze (Maze 1) over 15 trials to fully stabilize the behaviors (as reflected by a plateau in the time to reward), and then we blocked the shortest route by introducing a change point (Maze 2) prior to the 16th trial (Fig. 3A). Following this change, the average time to reward drastically increased, from 10 seconds to more than 1 minute (Fig. 3B), indicating that new effort and learning is required. Consistent with this, animals displayed significantly higher CA1 glucose drops in trial 16 (Maze 2) compared to trial 15 (Maze 1) (Fig. 3C), as revealed by mapping intracellular iGlucoSnFR2 signal dynamics onto the trajectories of the two trials (Fig. 3D and fig. S5). Moreover, within trial 16, when the mouse first passed through the corridor leading to the change point, which would have been interpreted as a “familiar” path, there was minimal glucose consumption compared to the return pass, at which point this same corridor would be interpreted as a “new” path (Fig. 3E, color tracks).

**Fig. 3.**
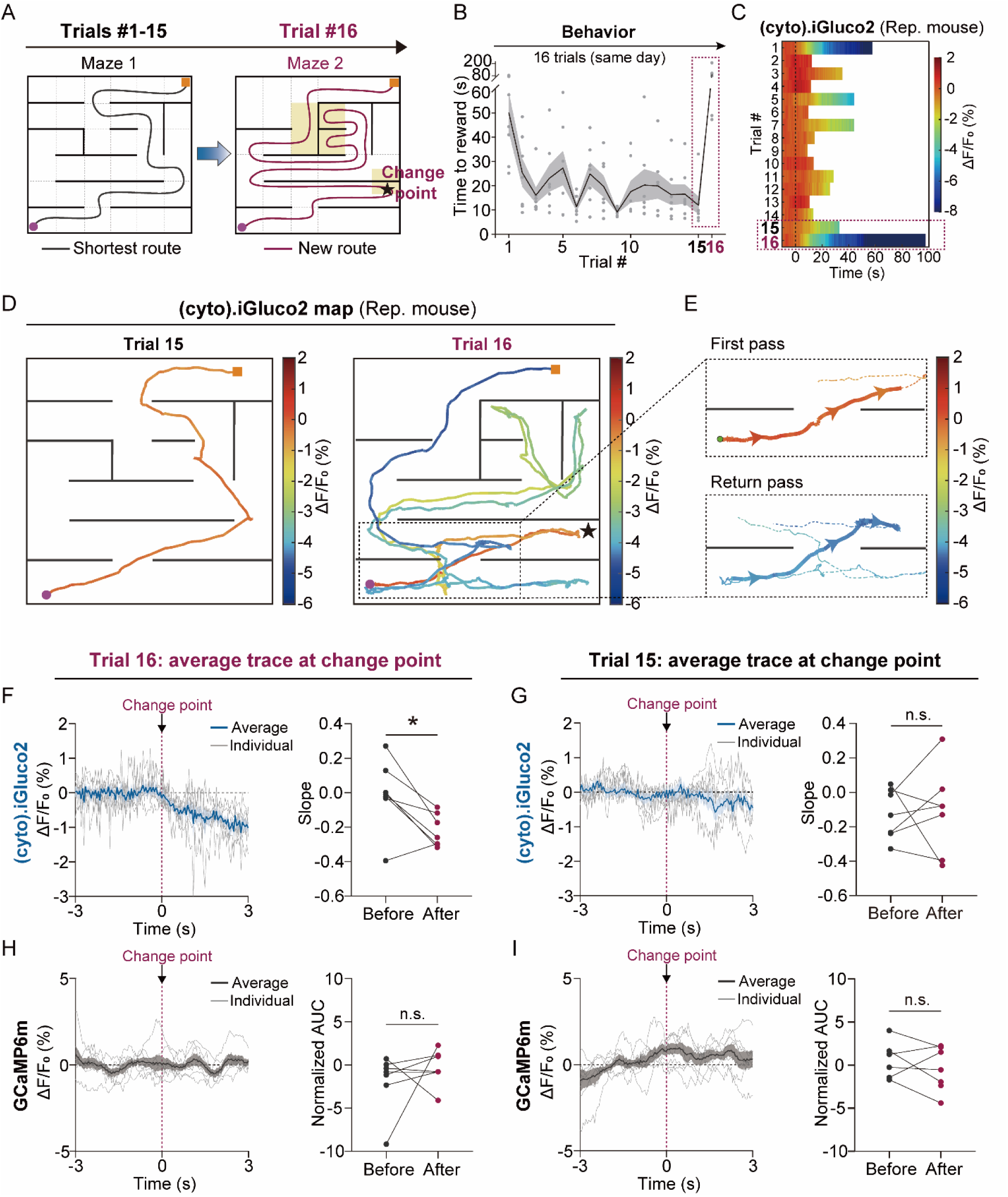
Glucose consumption rebounds during a renewed spatial navigation task. (A) Schematic illustrating the maze paradigm and the layout transition from maze 1 to maze 2. The black star indicates the change point in maze 2. (B) Maze navigation learning curve showing time to reward across trials 1–16 (n = 7 mice). (C) Trial onset–aligned (cyto).iGluco2 signals in CA1 neurons showing trial-by-trial dynamics across trials 1 to 16 in a representative mouse. (D) Mapping of (cyto).iGluco2 signal dynamics in CA1 neurons onto the trajectories from trials 15 and 16 in a representative mouse. (E) Enlarged view of the dashed-line region in trial 16 shown in (D). The upper trajectory shows the mouse initially passing through the corridor leading to the change point, whereas the lower trajectory shows the mouse traversing the same corridor after recognizing that the maze layout had changed. (F-G) Left: (cyto).iGluco2 signals surrounding the change point (±3 s) in trial 16 (F) and 15 (G). The blue trace represents the average signal across 7 mice, and the gray traces represent signals from individual mice. Right: quantification of the slope of the (cyto).iGluco2 signal before and after the change point. (H-I) Left: GCaMP6m signals surrounding the change point (±3 s) in trial 16 (H) and 15 (I). The black trace represents the average signal across 7 mice, and the gray traces represent signals from individual mice. Right: quantification of the normalized AUC of the GCaMP6m signal before and after the change point. Statistics were determined using a paired t test in (F to I). *P < 0.05.

Consistent with this observation, when we compared the intracellular glucose consumption before and after the point at which the maze was modified (change point), in trial 16, the “after” consumption was significantly higher than the “before” consumption, and there was no difference in consumption in trial 15 (before the modification was made) (Fig. 3, F and G). Together, these results indicate that it is the hippocampal representation, or subjective determination, of familiar versus new spatial information that dictates the energy consumption of the CA1 population, rather than the sensory information from a particular physical location.

Interestingly, in contrast to the changes in glucose consumption, re-learning did not significantly change bulk calcium dynamics (Fig. 3H and I). This dissociation suggests that CA1 energetic demand during spatial learning may not solely reflect changes in mean population activity. Instead, the increased glucose consumption may arise from metabolically demanding plasticity mechanisms required for rapid spatial map formation and updating (see discussion). Together, these findings suggest that CA1 glucose dynamics report the energetic cost of spatial mapping and remapping, revealing a form of learning-related energy demand that is not readily captured by bulk calcium activity alone.

### Glucose consumption of M1 neurons correlates with performance of motor learning

To determine whether post-learning glucose dynamics generalizes beyond rapid hippocampal spatial learning, we next examined a slower, cortex-dependent form of learning using the primary motor cortex (M1)-mediated single-pellet forelimb reaching task, an established paradigm for motor skill acquisition (*31–33*). In contrast to the maze task, which engages rapid CA1 map formation and updating, forelimb reach learning requires repeated practice over days and is associated with circuit remodeling in the motor cortex. This task also provides a more stereotyped behavioral output, minimizing potential confounding factors affecting systemic metabolism, such as variable trial durations, total locomotion and sensory integration across different maze locations.

We trained the mice for 7 sessions over 4 days using a modified single-hydrogel-pellet reaching task adapted from the classic single-pellet reaching protocol (*31, 32*), which increased their reward retrieval success from 20% to 50% (Fig. 4, A and B). AAV expressing either Syn-(cyto).iGlucoSnFR2 or Syn-GCaMP6m was injected bilaterally into the M1, and signals from the side contralateral to the preferred paw were analyzed to assess motor learning–induced changes (Fig. 4C and fig. S6A). During the early sessions (1 – 3), each successful reach was associated with higher glucose consumption (Fig. 4D). Neuronal glucose consumption of each successful reach significantly decreased as the animal progressed through motor learning session by session (Fig. 4D). As a negative control, we assessed CA1 glucose consumption, which was not altered during the motor training (Fig. 4E), again demonstrating that the post-learning change in energy consumption is task and circuit specific.

**Fig. 4.**
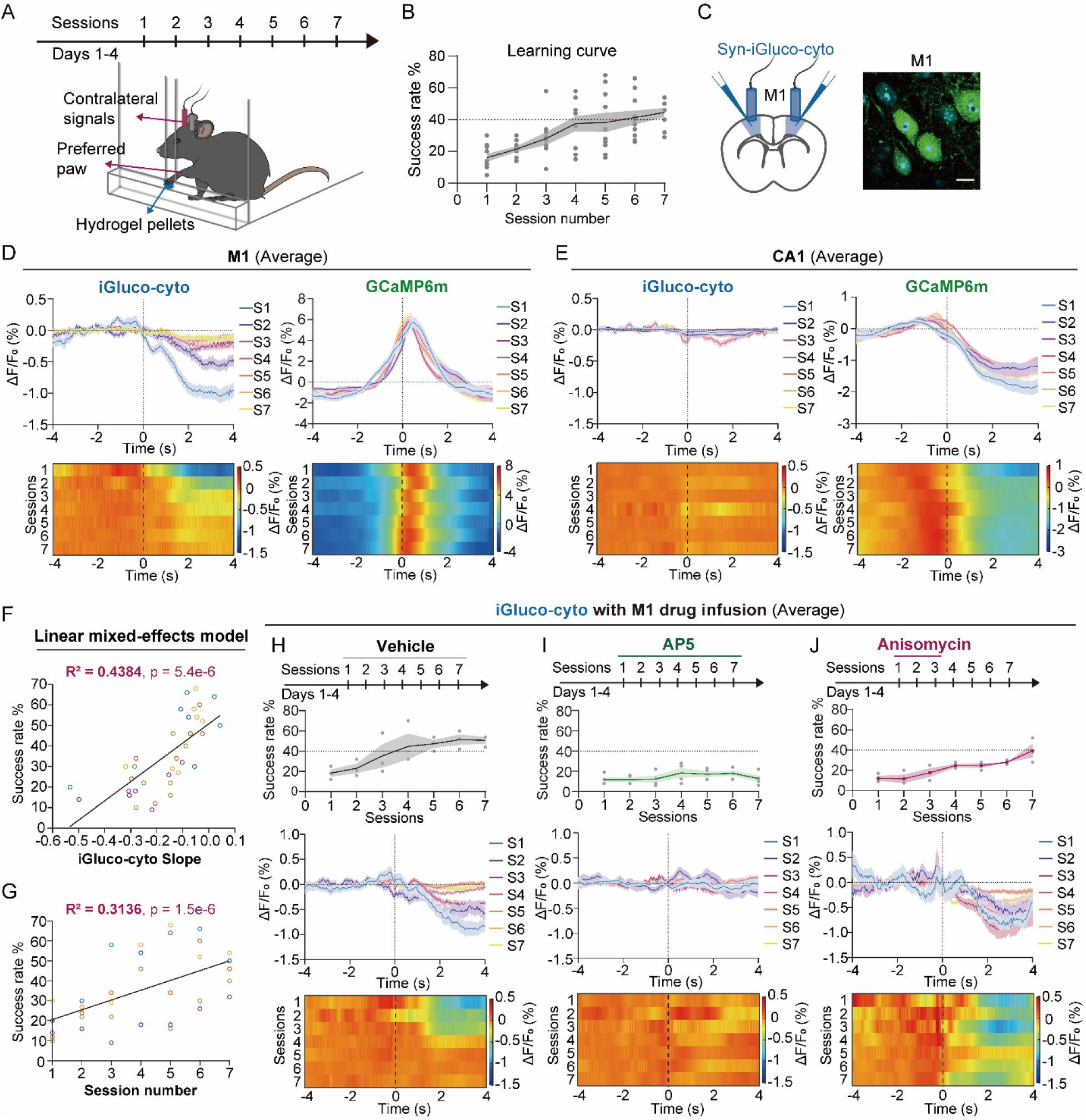
Glucose consumption of M1 neurons correlates with performance of motor learning. (A) Schematic of the motor learning paradigm. In the single-pellet forelimb reaching task, mice underwent 7 training sessions over 4 days. Water-deprived mice used their preferred forelimb to reach through a slit in the cage wall to grasp a hydrogel pellet placed on an external platform. Signals in M1 neurons from the side contralateral to the preferred paw were analyzed. (B) Motor learning curve for M1 (cyto).iGluco2 and GCaMP6m mice showing success rate across sessions 1–7 (n = 10 mice). (C) Left: schematic illustrating sensor injection and fiber placement in M1 cortex. AAV expressing either Syn-(cyto).iGluco2 or Syn-GCaMP6m was injected bilaterally into M1 cortex. Right: representative histology showing (cyto).iGluco2 expression in M1 neurons for fiber photometry recording (scale bar, 20 μm). (D) Average (cyto).iGluco2 and GCaMP6m signals in M1 neurons contralateral to the preferred paw across sessions 1–7, aligned to the onset of successful reaches in each session. Time 0 indicates the first reach toward the pellet (n = 5 mice for both (cyto).iGluco2 and GCaMP6m animals). (E) Average (cyto).iGluco2 and GCaMP6m signals in CA1 neurons across sessions 1–7, aligned to the onset of successful reaches in each session. (cyto).iGluco2 and GCaMP6m were expressed in the left and right CA1 of the same animals, respectively (n = 5 mice). (F–G) Correlation analysis of (cyto).iGluco2 slope versus success rate (F) and session number versus success rate (G) in M1 neurons during the motor learning task using a linear mixed-effects model. Data from the five mice are shown in five different colors. R^2^ represents the marginal R^2^ of the fixed effect ((cyto).iGluco2 slope or session number). (H–J) (cyto).iGluco2 signals in M1 neurons during the single-pellet forelimb reaching task with infusion of vehicle (H), AP5 (0.1 μg per site) (I), or anisomycin (25 μg per site) (J) into M1. Vehicle and AP5 were infused during each training session, and anisomycin was infused immediately after sessions 1–3. (cyto).iGluco2 expression, fiber placement, and drug infusion were all targeted to M1 contralateral to the preferred paw (n = 3 mice for the vehicle group, n = 4 for the AP5 group, and n = 3 for the anisomycin group).

As with CA1 neurons during spatial learning, once the motor skill has been learned and glucose consumption has stabilized at a low level, both the success rate and glucose consumption are insensitive to local glucose transporter inhibition (fig. S6, B to E). Consistent with a previous report (*34*), under this motor learning paradigm, the bulk M1 GCaMP6m activity did not significantly change as the mice mastered the motor skill (Fig. 4D). This may reflect the fact that motor learning reorganizes which M1 neurons are active and when, rather than altering the overall level of population activity (see discussion) (*35*).

Interestingly, the slope of iGlucoSnFR2 signals (the rate of intracellular glucose change across sessions) showed a significant positive correlation with trial success rate (R² = 0.4384, p = 5.4×10⁻⁶; Fig. 4F), indicating that mice with lower net glucose consumption achieved higher reaching success. Moreover, this correlation appeared stronger than that between success rate and session number alone (R² = 0.3136, p = 1.5×10⁻⁶; Fig. 4G), suggesting that the reduction in glucose consumption itself could be a more direct predictor of learning than elapsed training time.

### Different roles of NMDAR signaling and protein synthesis in post-learning glucose changes

The longer learning curve of the forelimb reaching learning paradigm provided an opportunity to use pharmacological manipulations to explore potential molecular mechanisms underlying energy optimization. Both NMDAR signaling (*36*) and new protein synthesis (*37*) play key roles in plasticity, learning and memory, and blockage of them has been shown to impair motor learning (*37–39*).

First, we infused the NMDA receptor antagonist AP5 into M1 to test how NMDAR-mediated mechanisms impact glucose consumption. Consistent with the literature that NMDAR is required for motor learning (*38, 39*), AP5 infusion robustly blocked behavioral performance (Fig. 4, H and I). Specifically, AP5 abolished the initial increase in glucose consumption in the first learning session, likely by blocking NMDAR-mediated calcium entry (Fig. 1M and fig. S2F). Accordingly, both glucose consumption and behavioral success rate remained low throughout subsequent training sessions (Fig. 4I).

Next, we examined how post-learning glucose consumption is affected by new protein synthesis. We adopted a protocol in which locally infusing anisomycin into M1 after each training session (sessions 1-3) robustly delayed motor learning (Fig. 4J) (*40*). With new protein synthesis blocked, the sharp glucose consumption induced by early training remained high and did not exhibit a post-learning reduction (Fig. 4J), as it did in the vehicle control group (Fig. 4H).

Together, these results suggest a two-step mechanism: an early NMDAR-dependent increase in Ca^2+^ entry that likely reflects the energy consumption of new plasticity events, followed by a subsequent consolidation phase in which protein synthesis plays a key role in remodeling the circuits to achieve a more energy-efficient processing protocol.

## Discussion

Using a genetically encoded glucose sensor in defined neuronal compartments, we directly measured the fuel cost of task-relevant neural processes in freely moving mice with sub-second resolution. Across multiple distinct learning paradigms, we found that learning was accompanied by a reduction in task-directed glucose consumption. This reduction was brain region-specific, closely tracked behavioral performance, rebounded during relearning, and depended on NMDA receptor signaling and protein synthesis. Together, these findings establish a quantitative and experimentally trackable link among energetic cost, neural activity, and learning.

The notion that the brain has evolved for energy efficiency is foundational in neuroscience. Principles such as sparse coding, minimized wiring, and signal-to-noise optimization are widely cited as evolutionary consequences of the metabolic constraints faced by neural systems (*1, 5, 8*). However, much of the evidence supporting this principle has come from theoretical models and from relatively static features of brain organization (*2–4, 41, 42*). By measuring *in vivo* fuel consumption within defined circuit nodes across diverse behaviors, our study provides experimental support for the energy-efficiency principle in action. These data suggest that, beyond the static architectural features of the brain that reflect long-term evolutionary optimization, the dynamics of learning and plasticity may themselves continuously and actively optimize metabolic cost.

At present, we do not yet have the ability to image glucose consumption at single-cell resolution in freely moving animals. Thus, beyond the observed requirements for NMDA receptor signaling and protein synthesis, the precise mechanisms by which learning alters glucose consumption remain an open question. Future development of cellular- or subcellular-resolution glucose imaging will be important for linking these metabolic changes to the extensive body of knowledge on neural plasticity. For example, in CA1, reductions in average firing rate (*30*), refinement of population tuning, and the formation of place fields (*43–45*) could each improve the signal-to-noise properties of neural coding and reduce the energetic cost of neural representation (*3, 8, 46*). Although our glucose measurements do not have the resolution of classical electrophysiological recordings, our GCaMP6m signals (Fig. 2, G and O) are broadly consistent with known patterns of neural activity. The rapid reduction in glucose consumption within one to two trials further suggests that behavioral time-scale synaptic plasticity (BTSP) (*47*) may contribute to this process. In motor learning, M1 neuronal activity becomes more stereotyped and more closely aligned with movement (*35, 48*), both of which could increase the SNR (*8*) and reduce the energetic cost of coding. At the synaptic level, learning reorganizes synapses by strengthening some connections while weakening others (*31, 49, 50*), and enables nonlinear dendritic computation (*32, 51–54*), processes that may reduce the number of spines or active synaptic elements required to support a given task and thereby lower energetic demand. These possible mechanisms in CA1 and M1 are not mutually exclusive and may operate in parallel. The potential contribution of astrocytes and other non-neuronal cells to learning-associated changes in glucose consumption also remains to be explored.

Although bulk glucose measurements lack cellular resolution, they are particularly meaningful for brain energetics because they capture physiologically relevant net fuel use. If the energetic cost measured in a single circuit is extrapolated to the scale of the whole brain, a brain that can perform the same task with less fuel would be expected to confer a selective advantage. Given this evolutionary pressure, and the consistency of post-learning reductions in energetic cost across behaviors, circuits, and timescales, we propose that energy-cost reduction may be a general outcome—and potentially an organizing objective—of learning at the cellular and circuit levels. We term this idea the “energy minimization hypothesis”: learning-induced plasticity moves neural network structure and dynamics toward configurations in which a neural task, such as transforming sensory input into an appropriate motor output sequence, can be accomplished at lower energetic cost.

In simplified terms, learning is a process through which an animal’s experience guides synaptic plasticity to generate new patterns of neural activity that improve future behavior (*55*). How these new patterns—such as memory ensembles (*56*), place fields (*45, 47, 57*), and activity trajectories (*35, 54, 58*)—are shaped by distinct plasticity mechanisms and instructive signals remains a central frontier in neuroscience. The energy minimization hypothesis does not aim to replace these mechanistic frameworks or directly answer all of these questions. Rather, it offers an orthogonal perspective on the trajectory of learning and plasticity. From this view, plasticity, broadly defined as changes in neural activity patterns and circuit organization, may guide circuits toward lower-energy configurations. This provides a simplified energetic lens for understanding brain computation and neural dynamics alongside the prevailing signal- and information-centered frameworks.

More broadly, this energetic view may also open new ways to think about intervention. Recent advances in drugs that potently modulate systemic and brain energy metabolism, including GLP-1 receptor agonists, have revealed unexpected effects on behavior and higher-order brain functions in both humans and animal models, extending beyond their established roles in food intake and metabolic control. These observations raise the possibility that targeting brain energetics may offer a new route to modulate neural function and, potentially, treat disorders in which learning, plasticity, or circuit dynamics are disrupted.

## Acknowledgements

We thank Chen Ran, Ardem Patapoutian, Anton Maximov, Xin Jin, Karl Deisseroth, Gulcin Pekkurnaz, Hollis Cline, and Ann Kennedy for discussion and feedback. We also thank all members of the Ye laboratory and the Dorris Neuroscience Center for their support and feedback.

## Author contributions

Conceptualization: LY; Methodology: LY, KX; Investigation: KX, FR, TQ, LS, BZ, XH; Reagent and Material: JSM; Funding acquisition: LY; Project administration: LY; Supervision: LY; Writing – original draft: LY, KX.

## Data, code, and materials availability

All data necessary to understand the conclusions of this study are available in the main text and supplementary materials.

## Materials and Methods

### Animals

All animal experiments were performed in accordance with the National Institutes of Health Guide for the Care and Use of Laboratory Animals and approved by Scripps Research Institutional Animal Care and Use Committee. Male C57BL/6J mice aged 8–10 weeks (Jackson Laboratory, stock no. 000664) were used for all experiments. Mice were group housed and maintained on a 12 h light: 12 h dark cycle with food and water ad libitum unless specified. Age-and body weight-matched mice were randomly assigned to experimental groups.

### Stereotaxic viral injection

AAV viruses, including AAV9-hSyn-(cyto).iGlucoSnFR2, AAV9-hSyn-(mem).iGlucoSnFR2, AAV9-hSyn-GCaMP6m (Addgene #100841), AAV9-hSyn-cpSFGFP, and AAV9-Gfap-(cyto).iGlucoSnFR2, were diluted in DPBS to a final titer of 5 × 10^12 vp/mL. A total volume of 300 nL of the diluted virus was injected into the primary visual cortex (lambda: AP, −0.5 mm; ML, 2.5 mm; DV, 1.25 mm), CA1 (bregma: AP, −1.8 mm; ML, 1.5 mm; DV, 1.35 mm), VMH (bregma: AP, −1.4 mm; ML, 0.5 mm; DV, 5.9 mm), or primary motor cortex (bregma: AP, 1.0 mm; ML, 1.5 mm; DV, 1.4 mm).

Mice were anesthetized with 1.5% isoflurane, and the skull was secured in a stereotaxic frame (David Kopf Instruments) and leveled using bregma and lambda as reference points. Virus was infused through a glass pipette at a rate of 100 nL/min using a nanoinjector (Nanoliter 2020 Injector, 300704, World Precision Instruments). After injection, the pipette was left in place for 5–10 min before being slowly withdrawn. For subsequent fiber implantation, either fiber-optic cannulas (400 μm core, 0.50 NA, RWD) or optical fiber multiple fluid injection cannulas (OmFC, Doric Lenses) were lowered to the target site and secured to the skull with dental cement (C&B-Metabond, Parkell). Mice were allowed to recover for at least 2 weeks before experiments.

### Fiber photometry

Fiber photometry was used to record GCaMP6m and iGlucoSnFR2 signals in freely behaving mice. Mice were connected to fiber-optic patch cables through implanted ferrules using ceramic mating sleeves. Patch cables consisted of single or double optical fibers with a 400 μm core diameter and 0.57 NA (Doric Lenses). The cables were coupled to 470 nm and 410 nm excitation light sources for signal and reference-channel recording, respectively. Before experiments, mice were habituated to the tethering system and remained tethered for at least 30 min before recordings began. The fiber photometry acquisition setup has been described previously (*59*). Fluorescence signals were acquired at 20 Hz. Reference-subtracted signals were expressed as ΔF/F₀ and aligned to behavioral events.

### Visual stimulation paradigm

Mice receiving bilateral AAV injections and optic fiber implants targeting the primary visual cortex were used in this paradigm. Two days prior to recording, animals were habituated to both the fiber-optic cable connection and the visual stimulation apparatus for 30 min/day. The apparatus comprised a transparent-walled chamber fitted with an internal light-blocking partition, with tablets displaying drifting sinusoidal gratings mounted along the exterior walls. The visual stimulus consisted of a full-field sinusoidal grating drifting perpendicular to its orientation at 2 Hz, and stimulus onset was triggered by raising the internal partition.

On the day of recording, mice first acclimated to the darkened chamber with the fiber-optic cable attached for 1 h before photometry acquisition commenced. In the standard stimulation protocol, ten 15-s visual epochs were presented over approximately 45 min, with 3–5 min inter-stimulus dark intervals between successive exposures.

### Maze navigation paradigm

All maze experiments were conducted using a Hebb-Williams maze (*29*) (MazeEngineers). The apparatus consisted of a 60 × 60 cm square arena with 10-cm-high walls. One corner served as the maze entrance, and the opposite corner contained a telemetry-controlled water reward device that delivered water for 2 s upon lick activation, followed by a 10-s no-delivery interval. The internal walls of the maze were movable, allowing different paths to be configured within the same arena. A camera mounted above the maze was used to record behavior throughout the experiments.

Preliminary training:the goal of preliminary training was to familiarize mice with the maze environment, the water port location, optic fiber cable connection, and handling, while establishing a reliable drinking habit confined to the maze. Successful training was defined as the ability to navigate directly to the water port upon maze entry with minimal fear or exploratory behavior, even as the internal maze configuration changed. Training was conducted over 4–5 days and comprised 8–10 sessions delivered at 12-h intervals. Throughout this period, water-restricted mice received water exclusively within the maze, and session duration was adjusted to ensure a minimum intake of 700 μL per session. Body weight was monitored daily, and the water restriction protocol was suspended if weight dropped below 80% of the initial weight. A simplified maze containing only two internal walls was used during preliminary training to facilitate water port discovery. Sessions progressed in a stepwise manner. In sessions 1 and 2, mice were placed in the simplified maze without fiber cable attachment for 15 min to explore the environment and learn the water port location. Sessions 3 and 4 were identical in structure but with the optic fiber cable connected, allowing habituation to the tethered condition. Beginning in sessions 5–10, a repeated handling procedure was introduced: upon each arrival at the water port, mice were permitted to drink for 10 s before being gently guided back to the entrance for the next trial. This cycle was repeated continuously over 15 min. Mice were considered ready for formal recording once they reached the water port and drank more than 20 times within 10 min across two consecutive sessions.

For formal recording, mice were tested in the maze configuration illustrated in Fig. 2B. Each session began with the mouse confined to the entrance alley for a 1-min baseline recording period. The door was then opened to initiate the first trial, during which the mouse freely navigated to the water port. Upon arrival, the mouse was allowed to drink for 10 s before being gently guided back to the entrance alley, where the door was closed for a 40-s inter-trial baseline period. The door was then reopened to begin the subsequent trial. This procedure was repeated for a total of six trials per session. Video recordings from the overhead camera were used for post-hoc behavioral annotation and trajectory tracking.

In the maze transition paradigm, mice were tested across two distinct maze configurations (Fig. 3A). Trials 1–15 were conducted in maze configuration 1, and trial 16 introduced maze configuration 2, allowing assessment of behavioral and neural responses to an unexpected change in the navigational environment.

### Motor learning paradigm

The single-pellet forelimb reaching task was used as the motor learning paradigm, performed as previously described (*31, 33*). We substituted food rewards with water rewards in the form of hydrogel pellets following 12 h of water restriction. to avoid the impact of hunger on systemic glucose level. This modification preserved the task structure and motivational drive while eliminating the metabolic interference of fluctuating systemic glucose on iGlucoSnFR2 signals recorded in M1. The training apparatus consisted of a Plexiglas chamber with a narrow vertical slit, in front of which a small platform was positioned to hold the hydrogel pellet within reaching distance.

Prior to formal training, all mice underwent habituation to the training chamber and were assessed for paw preference. During this assessment, hydrogel pellets were placed on the platform and mice were observed over 10 reaching attempts; the preferred paw was defined as the one used more frequently across attempts, or as the paw used in five consecutive reaches. Preliminary habituation then proceeded in a stepwise fashion: water-deprived mice were placed in the chamber for 15 min every 12 h and provided with 1 mL of hydrogel per session. The first two sessions were conducted without optic fiber cable connection, followed by two additional sessions with the cable attached, allowing progressive acclimation to the tethered recording condition.

Formal motor learning recordings spanned 4 days and comprised seven training sessions administered at 12-h intervals. Each session was terminated upon completion of 50 reaching attempts or after 20 min, whichever occurred first. Fiber photometry signals and behavioral video were recorded simultaneously throughout each session, with video timestamps used to align neural signals to discrete reaching events during post-hoc analysis. A reach was scored as successful when the mouse used its preferred paw to grasp the hydrogel pellet and bring it directly to its mouth without dropping it. Overall performance was quantified as the success rate, defined as the proportion of successful reaches relative to the total number of attempts per session.

### Local drug infusion using optical fiber multiple fluid injection cannulas

To enable local pharmacological manipulation of specific brain regions during fiber photometry recording, optical fiber multiple fluid injection cannulas (Doric Lenses) were implanted at sites of viral expression. Prior to each experiment, a fluid injector was inserted fully into the cannula guide tube and connected via polyethylene tubing to a 10-μL Hamilton syringe driven by a WPI SMARTouch controller.

Drug solutions were infused into the target region (V1, CA1, or M1) at a flow rate of 50–100 nL/min, with a total infusion volume of approximately 500 nL per site. The following compounds were used at the indicated doses: DRB18 (10 ng/site; MedChemExpress, HY-145963), muscimol (0.125 μg/site; Sigma-Aldrich, #M1523), AP5 (0.1 μg/site; Abcam, #ab120003), tetrodotoxin (TTX; 2 ng/site; Tocris, #1069), and anisomycin (25 μg/site; MedChemExpress, HY-18982).

### Histology and immunohistochemistry

Mice were deeply anesthetized with isoflurane and transcardially perfused with PBS followed by 4% PFA. Brains were then dissected and post-fixed overnight in 4% PFA at 4 °C. After fixation, brains were dehydrated in 30% sucrose for 24 h and embedded in OCT compound (Tissue-Tek O.C.T. Compound, Sakura). Coronal sections were prepared using a cryostat (FS800A, RWD). For histological analysis and immunostaining, 50-μm-thick sections were collected.

For immunostaining of the astrocyte marker S100B, free-floating sections were first blocked for 1 h at room temperature in blocking buffer consisting of 5% normal donkey serum in PBS containing 0.3% Triton X-100 (PBST). Sections were then incubated overnight at 4 °C with rabbit monoclonal anti-S100B antibody (E7C3A; Cell Signaling Technology, #90393) diluted 1:500 in blocking buffer. After primary antibody incubation, sections were washed three times in PBST for 15 min each and then incubated for 1 h at room temperature with donkey anti-rabbit IgG secondary antibody conjugated to Alexa Fluor 647, diluted 1:500 in blocking buffer. Sections were subsequently washed three times in PBST for 15 min each, stained with DAPI (5 nM in PBS) for 30 min, and mounted in Fluoromount-G (Electron Microscopy Sciences) for imaging with a confocal microscope (FV4000, Olympus).

### Data analysis and quantification

Fiber Photometry Preprocessing: for all paradigms, raw fluorescence signals first reference-subtracted and subsequently normalized as ΔF/F₀ using the entire recording prior to further analysis. Trial- or reach-aligned ΔF/F₀ traces were computed relative to a baseline period defined for each paradigm as described below and were used for heatmap visualization and time-course comparisons throughout.

For fiber photometry data analysis in the visual stimulation paradigm shown in Fig. 1, traces were aligned to the onset of visual stimulation and averaged across stimulation trials. For each visual stimulation, the mean signal during the 30-s period preceding the 15-s visual stimulation epoch was used as the baseline to zero-normalize the onset trace. To compare (cyto).iGlucoSnFR2, (mem).iGlucoSnFR2, and cpSFGFP signals, the normalized area under the curve (AUC) during the 15-s visual stimulation period was quantified and used for statistical analysis.

For fiber photometry data analysis in the maze navigation paradigm shown in Fig. 2 and 3, preprocessed signals were segmented on a trial-by-trial basis according to behavioral annotations. Trial onset was defined as the moment the mouse entered the maze (time 0), and trial end as the moment of water port arrival. For each trial, the mean signal of the 20-s period preceding maze entry was used as the baseline to zero the signal. For cross-trial comparisons, GCaMP6m, (cyto).iGlucoSnFR2 and (mem).iGlucoSnFR2 signals were quantified as the normalized area under the curve (AUC) computed from trial onset to trial end. To account for variability in trial duration, AUC values were normalized by trial length.

Mouse trajectories were extracted from overhead video recordings using DeepLabCut (*60*). The body center was tracked frame-by-frame, and the resulting x–y coordinates were aligned to behavioral annotations. Trial-specific trajectories from maze entry to water port arrival were then reconstructed to visualize each animal’s navigation path.

For fiber photometry data analysis in the maze transition paradigm shown in Fig. 3, trial-aligned GCaMP6m and (cyto).iGlucoSnFR2 ΔF/F₀ signals were mapped onto the reconstructed mouse trajectories to visualize the spatial and temporal dynamics of neural and metabolic signals along the navigation path, enabling direct comparison between trial 15 (familiar configuration) and trial 16 (novel configuration). For the change-point analysis in Fig. 3, signals immediately before and after the change point were extracted using windows of up to 3 s with window boundaries constrained by the start and end of each trial. (cyto).iGlucoSnFR2 responses were quantified as the slope of the ΔF/F₀ trace obtained by linear regression within each pre- or post-change-point window, whereas GCaMP6m responses were quantified using the normalized AUC over the corresponding window (re-zeroed to the starting value of the window).

For fiber photometry analysis in the motor learning paradigm, signals recorded from M1 neurons contralateral to the preferred paw were used for analysis. Fluorescence signals were reference-subtracted and segmented reach-by-reach according to behavioral annotations. Analyses were restricted to successful reaching events, with time 0 defined as the moment the mouse first placed its paw on the hydrogel pellet. Successful reach-aligned ΔF/F₀ traces were used to generate session-averaged time-course plots and heatmaps. The relationship between motor performance and (cyto).iGlucoSnFR2 dynamics was assessed at the session level: for each mouse and session, individual reach-aligned (cyto).iGlucoSnFR2 ΔF/F₀ traces were converted to percent ΔF/F₀ and averaged into a single session-averaged trace. The glucose response was then quantified as the slope of this averaged trace from 0 to 2 s after movement onset using linear regression. For each mouse and session, the (cyto).iGlucoSnFR2 slope was paired with the corresponding behavioral success rate, and two separate linear mixed-effects models were fitted: one testing the effect of session number on success rate, and the other testing the effect of (cyto).iGlucoSnFR2 slope on success rate. In both models, the predictor (session number or glucose slope) was included as a fixed effect, with mouse identity as a random intercept to account for repeated measurements across sessions within the same animal. Marginal R^2^ of the fixed effects was used.

### Statistics

Statistical analyses were performed using Prism 10 (GraphPad Software) for paired and unpaired t tests. The specific statistical tests used and the number of biological replicates are indicated in the corresponding figure legends. A P value of less than 0.05 was considered statistically significant.

---

**Fig. S1.**
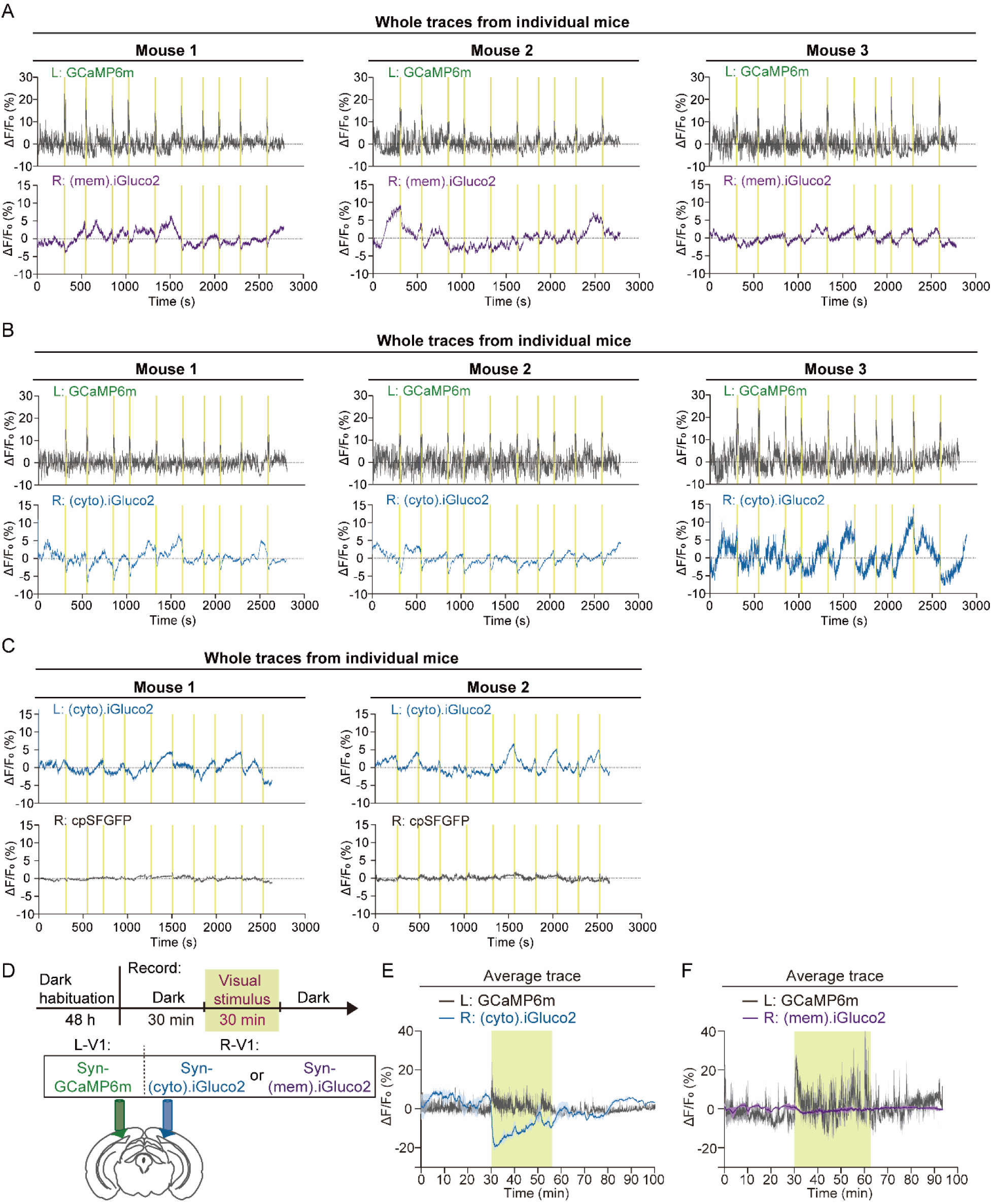
Visual stimulation reduces glucose levels in primary visual cortex neurons. (A) Individual mouse traces showing GCaMP6m and (mem).iGluco2 signals recorded from the left (L, in green) and right (R, in purple) V1 neurons of the same animal. (B) Individual mouse traces showing GCaMP6m and (cyto).iGluco2 signals recorded from the left and right V1 neurons of the same animal. (C) Individual mouse traces showing (cyto).iGluco2 and cpSFGFP signals recorded from the left and right V1 neurons of the same animal. (D) Schematic showing the long-term 30-min visual stimulation paradigm, sensor injection, and fiber placement for the experiments in E and F. (E) Average fiber photometry traces showing GCaMP6m signals in the left V1 and (cyto).iGluco2 signals in the right V1 of the same animal during visual stimulation (n = 3 mice). (F) Average fiber photometry traces showing GCaMP6m signals in the left V1 and (mem).iGluco2 signals in the right V1 of the same animal during visual stimulation (n = 3 mice).

**Fig. S2.**
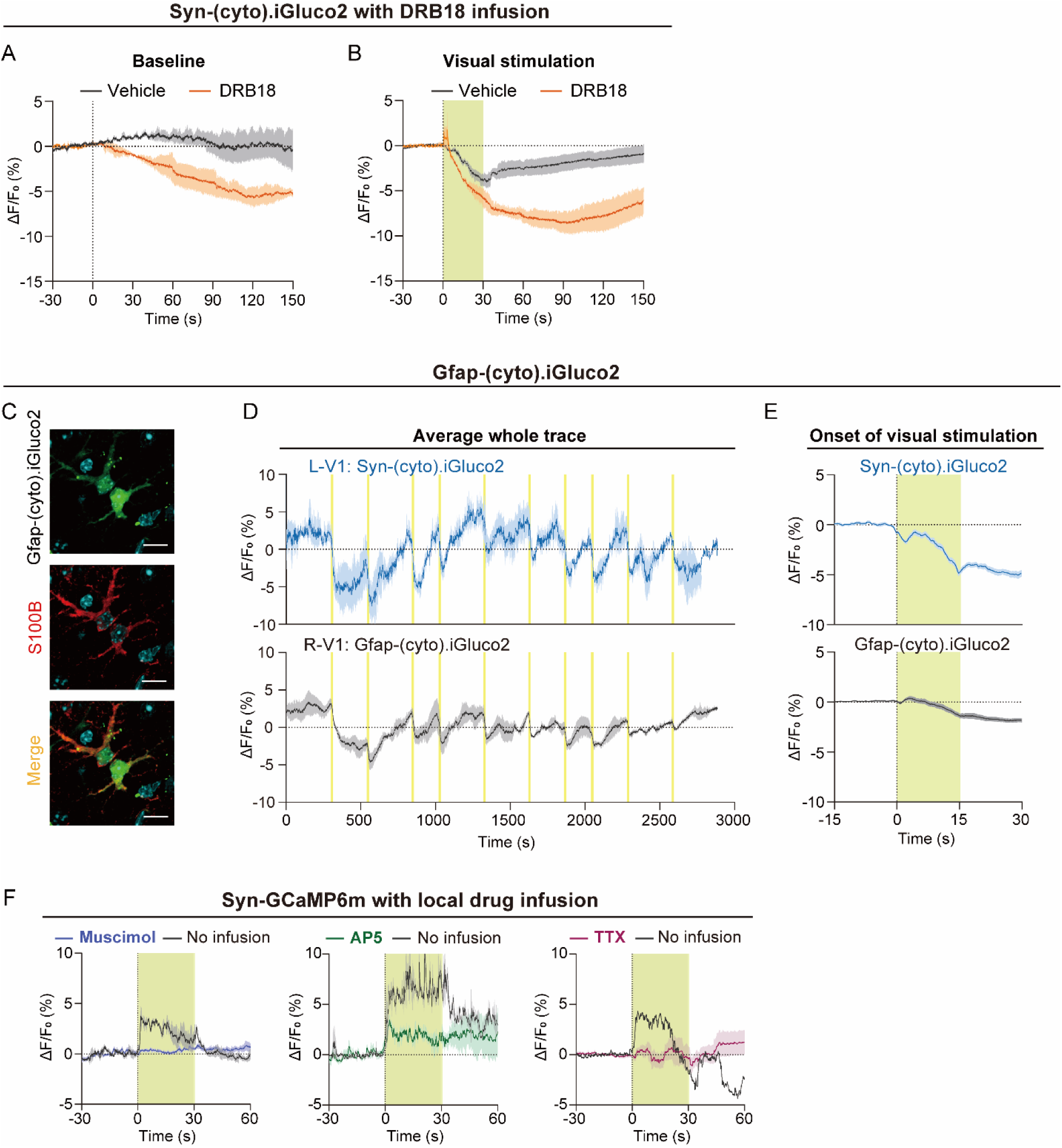
Visual stimulation-induced neuronal glucose drop is driven by neuronal activity. (A-B) (cyto).iGluco2 signals in V1 neurons following vehicle or pan-GLUT transporter inhibitor (DRB18) infusion under baseline (A) and visual stimulation (B) conditions. (C) Histology showing colocalization of Gfap-(cyto).iGluco2 expression with the astrocyte marker S100B in V1 cortex. (D) Average fiber photometry traces showing Syn-(cyto).iGluco2 signals in the left V1 (upper) and Gfap-(cyto).iGluco2 signals in the right V1 (lower) of the same animal during visual stimulation (n = 3 mice). (E) Signal changes aligned to the onset of visual stimulation (n = 3 mice) (F) GCaMP6m signal changes aligned to visual stimulation onset with and without local drug infusion. A GABA_A_ receptor agonist (muscimol), an NMDAR antagonist (AP5), or a sodium channel blocker (TTX) was infused into V1 through the cannula.

**Fig. S3.**
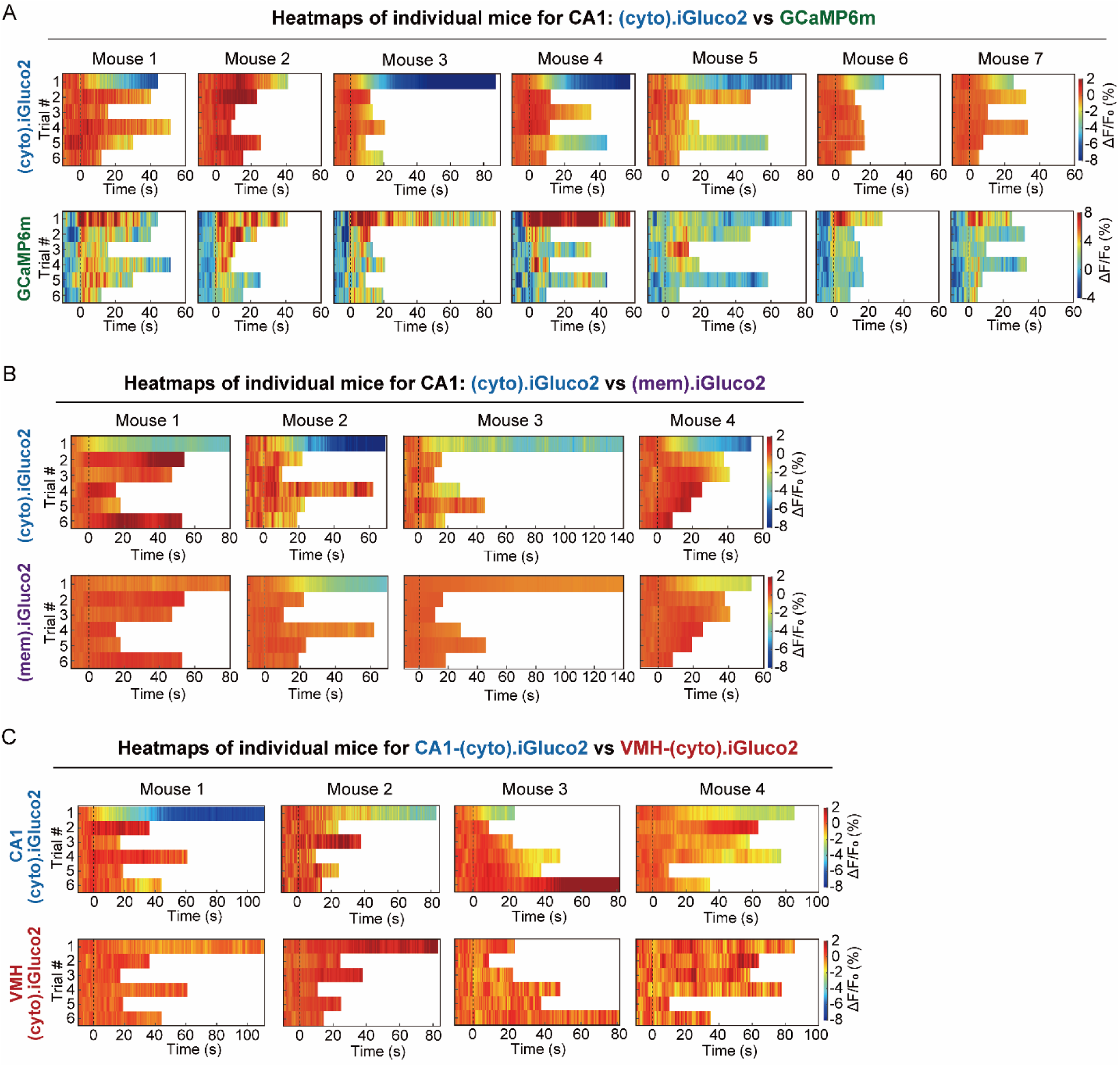
Spatial navigation learning leads to a reduction in glucose consumption in CA1 neurons. (A) Heatmaps from individual mice showing (cyto).iGluco2 and GCaMP6m signals in CA1 neurons across trials 1–6 during the maze task in Fig. 2 F-H. (B) Heatmaps from individual mice showing (cyto).iGluco2 and (mem).iGluco2 signals in CA1 neurons across trials 1–6 during the maze task in Fig. 2 I-K. (C) Heatmaps from individual mice showing (cyto).iGluco2 signals in CA1 and VMH neurons across trials 1–6 during the maze task in Fig. 2 L-N.

**Fig. S4.**
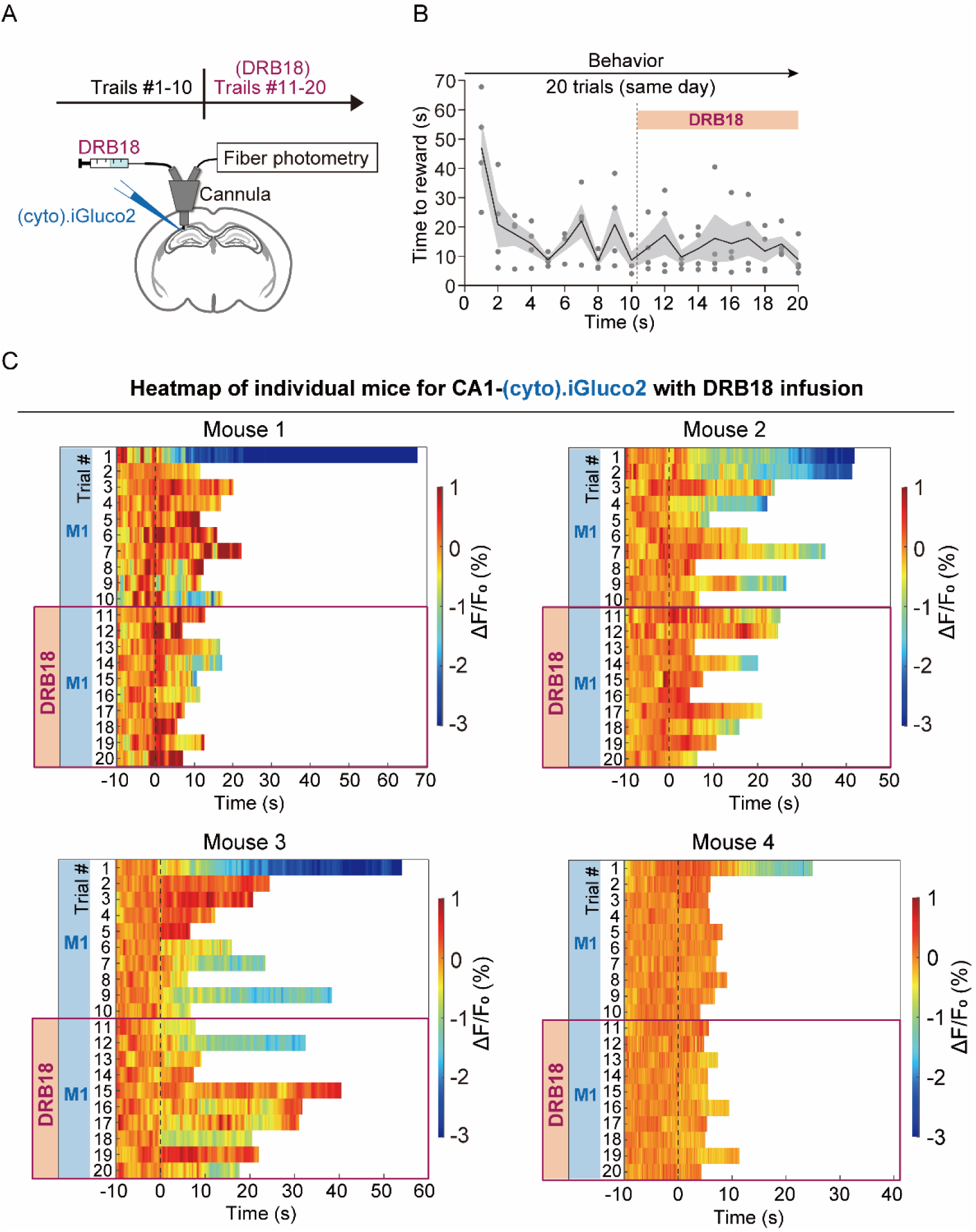
CA1 neurons maintain low glucose consumption in the familiar maze. (A) Schematic showing sensor injection and cannula placement for local drug infusion in CA1 during fiber photometry recording. (B) Maze navigation learning curve showing the time to reward across trials 1–20 in the maze task following DRB18 infusion (n = 4 mice). (C) Heatmaps from individual mice showing (cyto).iGluco2 signals in CA1 neurons across trials 1–20 in the maze task following DRB18 infusion.

**Fig. S5.**
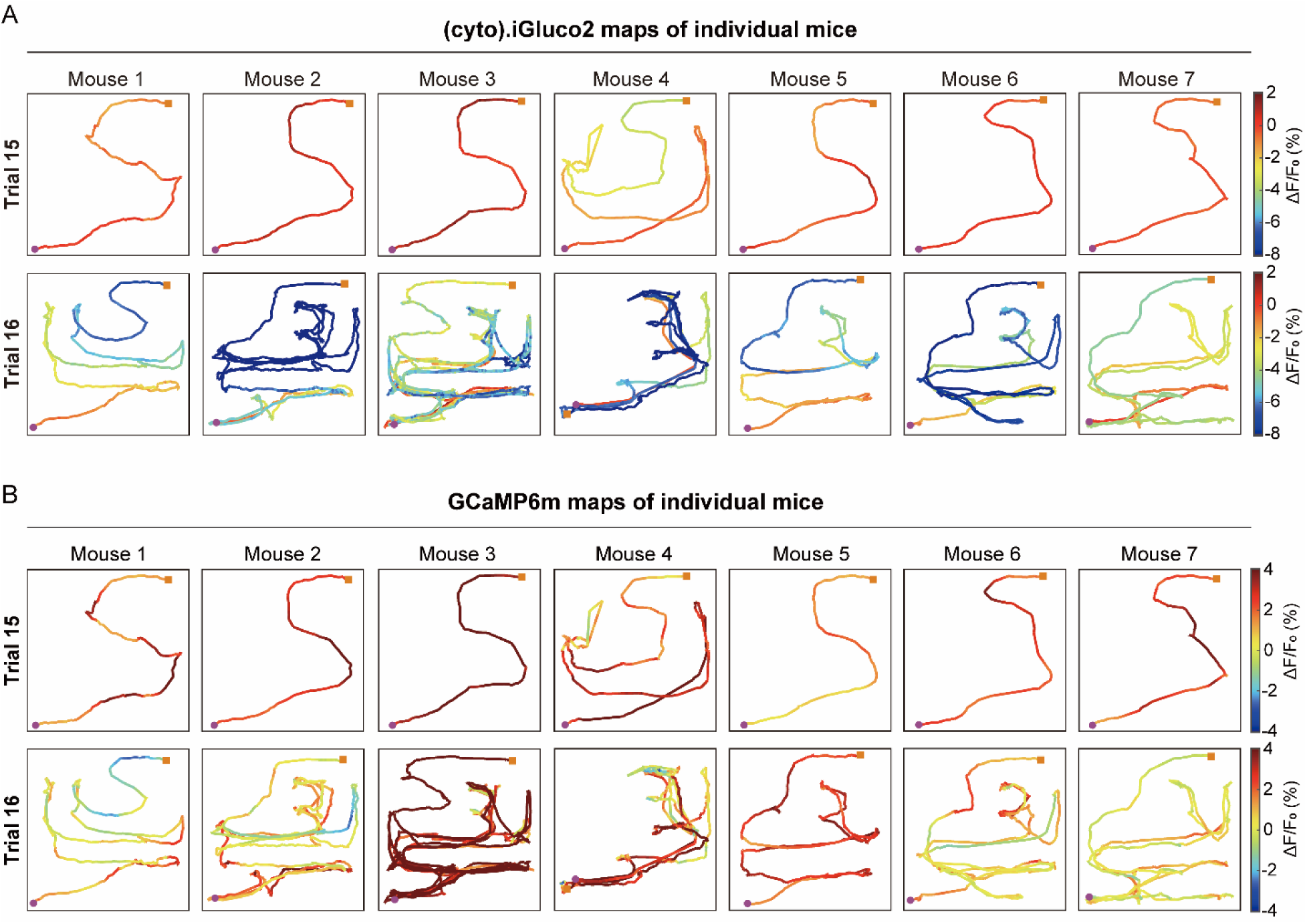
Intracellular glucose and calcium dynamics in CA1 neurons during maze transition. (A) Maps of (cyto).iGluco2 signal dynamics in CA1 neurons during trials 15 and 16 for individual mice. (B) Maps of GCaMP6m signal dynamics in CA1 neurons during trials 15 and 16 for individual mice.

**Fig. S6.**
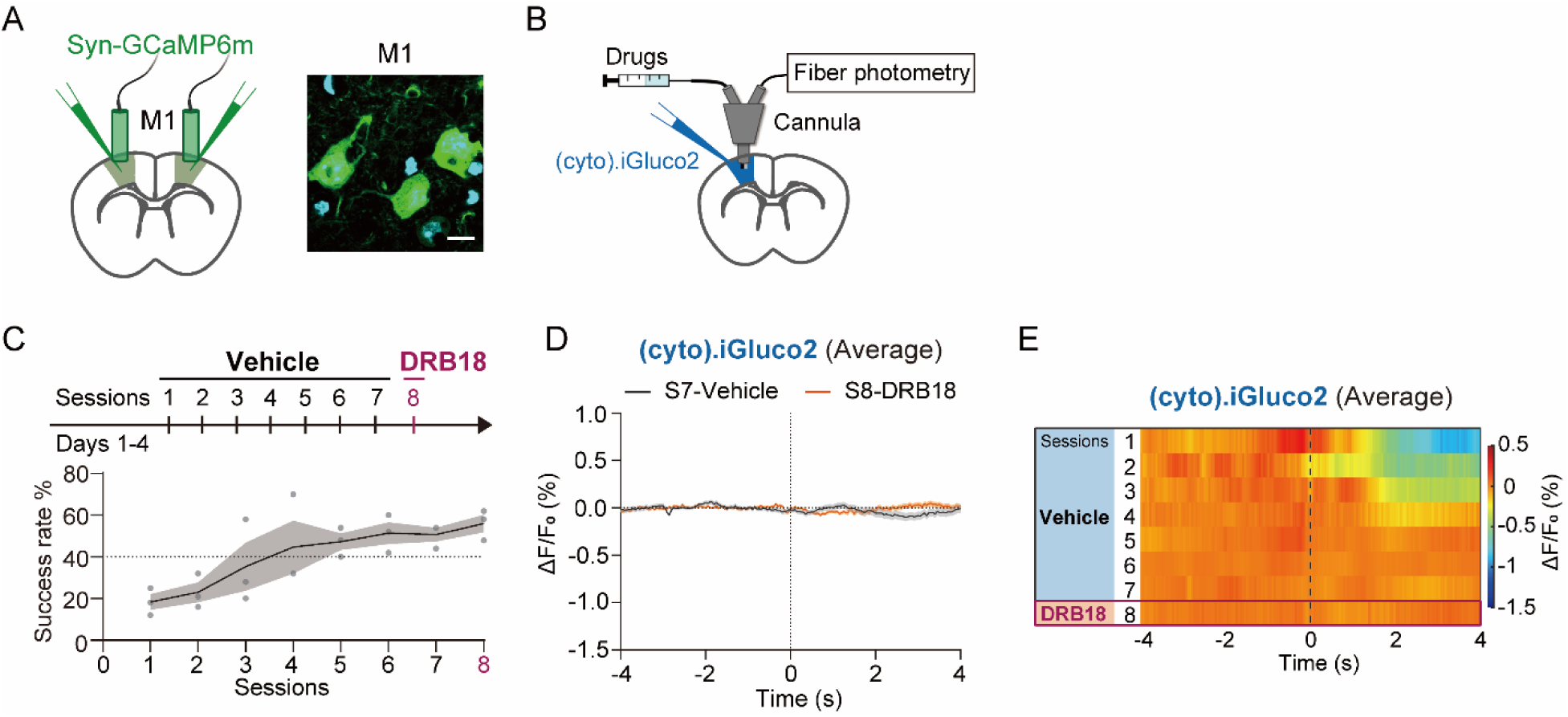
M1 neurons maintain stably low glucose consumption after motor skill acquisition. (A) Schematic illustrating sensor injection and fiber placement in M1 cortex (left). Representative histology showing GCaMP6m expression in M1 neurons for fiber photometry recording (right, scale bar, 20 μm). (B) Schematic showing sensor injection and cannula placement for local drug infusion in M1 during fiber photometry recording. (C) Motor learning curve for M1 (cyto).iGluco2 mice showing success rate across sessions 1–8, with vehicle infused during sessions 1–7 and DRB18 infused in session 8 (n = 3 mice). (D-E) Average (cyto).iGluco2 signal in M1 neurons aligned to the onset of successful reaches in session 7 and 8 (n = 3 mice).

